# Tree-to-cycle transition scale characterizes trade-off between dissipation and construction cost in adaptive transport networks

**DOI:** 10.64898/2026.09.02.748799

**Authors:** Justina Stark, Christophe Godin

## Abstract

Biological transport networks range from tree-like to highly reticulated architectures, which have been proposed to reflect different balances between viscous dissipation and metabolic cost. This balance cannot currently be inferred from structure: available descriptors either discard edge width entirely or preserve it only as a hierarchical decomposition that has not been mapped to the dissipation–cost trade-off. Here we introduce the tree-to-cycle transition scale, a single number given by the radius of the thickest edge outside the maximum spanning tree, which we interpret as the highest cost a network accepts for redundancy. We compute it for networks adapting to spatially correlated load fluctuations, whose correlation length sets their position on the Pareto front. The transition scale decreases linearly with dissipation and increases linearly with metabolic cost. Its scatter is largest in the regime where some networks end up off the front, and adding a growth term removes the scatter but splits the correlation into two branches. A single quantity read off the static network architecture thus characterizes where a network sits on the Pareto front, and is sensitive to the trajectory the network took through the optimization landscape. This suggests a route to comparing observed networks, for example leaf venations across species or growth conditions, by the trade-off they realize.

## 1 Introduction

Biological transport networks, such as vasculatures in animals and plants, provide vital routes for tissue supply and clearance. These networks are required to perform multiple functional objectives, such as robustly meeting the tissue demands while minimizing the cost of transport and network construction (Kramer et al. 2023). As these objectives compete (e.g., enhancing robustness by adding connections also increases the construction cost), there exists no single optimal solution to this optimization problem, but a set of non-dominated, i.e., equally valid trade-off solutions. These trade-off solutions form a sub-space in the multi-objective space, called the *Pareto front*, along which no objective can be improved without worsening another (Farnsworth et al. 1995). The position of a solution along the Pareto front depends on how the trade-off is actually balanced.

Different ways to balance the competing objectives have been proposed to underlie the variety in network architectures found in nature, which ranges from tree-like to highly cyclic structures (Blonder et al. 2020; Matos et al. 2024; Ronellenfitsch et al. 2019); e.g., animal arteries and plant stems typically adopt a fractal, tree-like architecture that minimizes transport and material cost, whereas animal capillaries and leaf venation gain robustness against damage and environmental fluctuations through cyclic components (Corson 2010; Katifori et al. 2010; Waszkiewicz et al. 2024). These cyclic components are usually formed at a smaller spatial scale by thinner network edges (e.g., leaf minor veins), while thicker edges of the same organ remain tree-like at larger scale (e.g., leaf major veins). This suggests that a scale exists, related to the thickness of the network edges (Kaiser et al. 2020), at which a redundant connection’s additional cost is outweighed by its benefits.

Theoretical models have successfully predicted the emergence of such architectures as different trade-off balances (Ronellenfitsch et al. 2019). The inverse, however — inferring a network’s trade-off balance and performance from its observed architecture — remains unresolved (Matos et al. 2024); this is because characterization of variable and diverse networks requires descriptors that map complex architectures to a scalar value, and standard network descriptors used for this purpose, such as degree distribution, density, and cycle count (Barthélemy 2011), are topology-based and therefore do not preserve edge width, which, together with topology, determines a network’s dissipation and metabolic cost and thus its trade-off balance. Descriptors that do retain edge weights, such as the hierarchical loop decomposition (Katifori et al. 2012) and its extension to non-planar networks (Modes et al. 2016), instead map the network to a nested tree whose branches record the order in which cycles merge as the weakest edges are removed. These decompositions characterize hierarchical organization, but no map has been established from them to a network’s position in the dissipation versus metabolic cost plane. Existing descriptors can therefore not quantify how the trade-off is balanced by different network architectures and predict how this affects network function. This limitation prevents the use of network architecture as a trait index for inferring functional performance and adaptation, e.g., in ecology or crop engineering.

Here, we use the scale at which a network transitions from a tree-like to a cyclic structure as a descriptor linking network architecture to functional trade-off. We find that this tree-to-cycle transition scale (1) correlates with network position in the multi-objective space, and (2) is sensitive to the trajectory along which the networks were adapted.

## 2 Supply network modeling

We consider simple supply networks modeled as graphs with a set of nodes *N* connected by a set of undirected edges *E*, where each pair of adjacent nodes *i* and *j* contributes a single element (*i, j*) ∈ *E* and the ordering within the pair fixes only the reference direction of the flow. Each node carries a potential (e.g., pressure, concentration, or voltage), and if a potential difference exists between adjacent nodes *i* and *j*, a volumetric flow rate *q*_*ij*_ (volume/-time) is induced along the edge (*i, j*) (Marbach et al. 2023), i.e.:

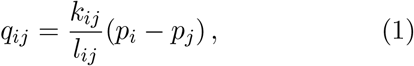

with 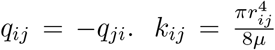 is the conductivity of an edge with radius *r*_*ij*_ and fluid viscosity *µ* and *l*_*ij*_ is the edge length. Flow continuity at the nodes according to Kirchhoff’s current law is satisfied by imposing Neumann boundary conditions such that incoming and outgoing flows at a node *i* sum to the net source or sink strength *s*_*i*_ at that node,

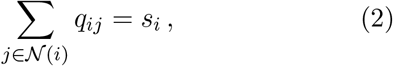

here *N*(*i*) are the adjacent nodes connected to *I* and

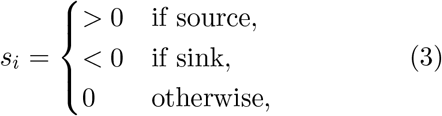

and ∑_*i*∈*N*_ *s*_*i*_ = 0. Fixing ***s*** = {*s*_*i*_} and ***k*** = {*k*_*ij*_}, the node pressures ***p*** = {*p*_*i*_} are obtained by substituting Eq. (1) into Eq. (2) and then solving ℒ***p*** = ***s*** for ***p***, where ℒ is the |*N* |×|*N* | weighted graph Laplacian, i.e., ℒ_*ii*_ = _*j*∈*N* (*i*)_ *k*_*ij*_*/l*_*ij*_ and ℒ_*ij*_ = −*k*_*ij*_*/l*_*ij*_ if *j* ∈ *N*(*i*), otherwise ℒ_*ij*_ = 0. Using these node pressures, Eq. (1) then yields the networks flow vector ***q*** = {*q*_*ij*_}.

In many animal systems, vascular branching angles and radii have been theoretically demonstrated to be consistent with cost optimization principles, minimizing the combined energy cost of transporting the blood and maintaining vessel and blood tissues (Cecil D Murray 1926a,b; Zamir 1976, 1977). The work to maintain a flow, e.g., in the case of fluid flow to overcome viscous resistance (Zamir 1977), corresponds to the network’s dissipation *D* per unit time (Corson 2010; Katifori et al. 2010):

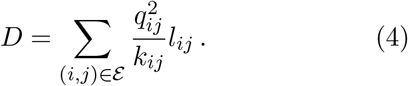

This means that the energy cost associated with transport can be reduced by increasing the conductivity of edges with large *q*, e.g., by dilating a vein that carries larger flows. However, increasing the conductance causes an increase in the metabolic cost *M* of constructing and maintaining the network’s components (e.g., tubes, blood volume) (Bohn et al. 2007; Hu et al. 2013; Ronellenfitsch et al. 2019):

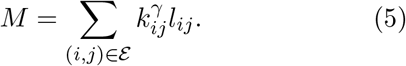

*M* scales linearly with *l*_*ij*_ because the amount of required material of a vein is always proportional to its length. While the length is imposed by the system size, the organism can influence *M* by adapting the conductivity (via the radius of the edge segment). The scaling of *M* with the conductivity, *γ*, depends on the network type: because *k* ∝ *r*^4^, *γ* = 0.25 if *M* scales with the mass of a thin-walled pipe (mass_pipe_ ∝ *r*). *γ* = 0.5 if *M* scales with the fluid volume (*V* ∝ *r*^2^), in the case of which Eq. (5) equals the cost of blood. *γ* = 1.0 if *M* scales with the mass of bundles of identical tubes (*M*_bundle_ ∝ *n*_tubes_ × *k*_singletube_). The architecture of animal or plant vasculatures has been shown to be consistent with 0.5 ≤ *γ <* 1.0 (Hu et al. 2013; Kaiser et al. 2020; Klemm et al. 2023; Ronellenfitsch et al. 2019). For 0 *< γ <* 1, the network minimizing *E* is a tree (i.e., cycle-free) (Banavar et al. 2000; Bohn et al. 2007; Burger et al. 2019), whereas for *γ >* 1 adding edges is cheaper than increasing *k*, and the network minimizing *E* is the full lattice (Bohn et al. 2007; Hu et al. 2013). *γ* thus parametrizes the optimal network’s phase transition from a hierarchical tree architecture to a network with cycles (Bohn et al. 2007). This phase transition is, however, shifted towards *γ <* 1 if flows fluctuate (Corson 2010) or if edges are randomly damaged (Katifori et al. 2010). Networks can also dynamically optimize their architecture as they adapt to the transported flows; in plants, venation networks are patterned by a hormone which enhances its own transport by a flux-induced feedback (Rolland-Lagan et al. 2005), and in animals, vasculatures remodel to a flow-induced wall shear stress (Espina et al. 2023; Fisher et al. 2001). Motivated by the latter, the dynamic vessel adaptation equation has been derived (Hu et al. 2013):

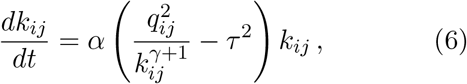

which satisfies a gradient-descent-like dynamics on *E* (adaptation time *t*, rate constant *α*), i.e., *dE/dt* ≤ 0 along its trajectories, driving the network toward a (locally) energy-minimizing configuration (Hu et al. 2013). Eq. (6) has been constructed such that, for *γ* = 0.5 (blood cost), edge conductivities are driven towards a state where the mechanical driving stimulus from the flow 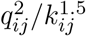 equals the squared target wall shear stress *τ* ^2^. This means that close to steady state, Eq. (6) converges to the experimentally observed wall shear stress adaptation (Hu et al. 2013).

Consistent with the previously discussed global network optimizations (Bohn et al. 2007; Corson 2010; Katifori et al. 2010), the steady-state solution of Eq. (6) is a tree for *γ <* 1.0 and a mesh-like network for *γ >* 1.0, with this phase transition of cyclic components shifted to *γ <* 1.0 if flows fluctuate (Hu et al. 2013).

Such fluctuating flows are modeled based on the assumption that adaptation is much slower than the timescale of fluctuations, so that the effective adaptation stimulus is the second raw moment of the flow (Lu et al. 2021) ^1^, 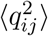, averaged over an ensemble of source/sink configurations (Hu et al. 2013; Ronel-lenfitsch et al. 2019). The choice of this ensemble determines which architectures emerge.

Ronellenfitsch et al. (2019) describe adaptation to a uniformly weighted ensemble of spatially correlated sink patterns with a fixed source. Following this model, we consider a system of one source node, indexed *l* = 0, which carries a fixed inflow of +1 in every microstate, and |*N* | − 1 sink nodes over which the outflow is distributed. A fluctuation centered on node *m* defines one microstate, where *m* runs over all |*N* | nodes, so that the source-sink vector 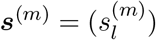 is:

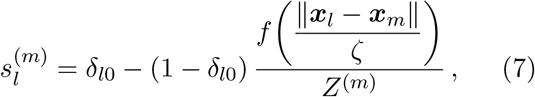

where ∥***x***_*l*_ − ***x***_*m*_∥ is the Euclidean distance between node *l* and the fluctuation center *m*, and *δ* is the Kronecker delta. *f* is the kernel function that determines how a fluctuation at *m* influences the sink strength at *l*. The kernel function *f* has to satisfy *f* (0) = 1 and *f* (∞) = 0. We use the Gaussian 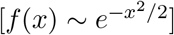, but other kernel functions have been shown to lead to similar results (Ronellenfitsch et al. 2019). The kernel width is scaled by *ζ* so that sinks are highly correlated for large *ζ* and uncorrelated if *ζ* → 0. The total outflow of each microstate is normalized to −1 by the sum of the kernel over all sinks, matching the inflow +1 at the source,

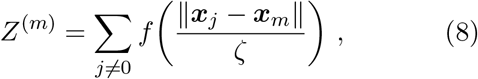

so that 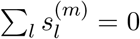 for every *m*. Note that the normalizing sum in Eq. (8) excludes the source node, whereas the microstate index *m* runs over all *N* nodes, i.e., the ensemble includes the state in which the sink is concentrated around the source. The average adaptation stimulus over all *N* microstates is

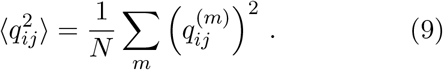

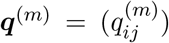 is the flow vector of microstate *m*. A limitation of optimizing networks by Eq. (6) is that it can get stuck in local minima of *E*. It has been proposed to reduce this risk by adding a flow-independent term *ce*^−*λt*^ to the adaptation dynamics, which drives the network along a different trajectory through the energy landscape. This has been interpreted as modeling the effect of tissue growth, and it has been shown to improve the optimality of the final network (Ronellenfitsch et al. 2016). One potential reason for this improved optimality is that the flow-independent term adds the same increment to every edge and thus keeps thin edges from being pruned early; the network commits to a topology only once the flow-dependent term dominates, which may prevent trapping in local minima during the early adaptation phase. The general dimensionless adaptation equation is (adapted from Ronellenfitsch et al. 2019):

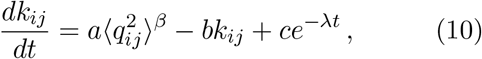

where each term is now weighted individually by adaptation parameters *a, b*, and *c*, and *β* = 1*/*(*γ*+1), i.e., *β* = 2*/*3 has the same steady state as the shear stress adaptation in Eq. (6). Eq. (7) – (10) enable sampling the space of Pareto-efficient networks whose balances of *D* and *M* are controlled by the correlation length scale *ζ*. In particular, if *ζ* is large, sink fluctuations overlap, and the variance between microstates is small, so that the effective stimulus is similar to that of steady flow, resulting in tree-like networks; whereas if *ζ* is small, sink fluctuations overlap less, the variance is larger, and resulting networks have cycles, i.e., they are reticulated. The tree-like networks lie at the end of the Pareto front minimizing *M* at larger *D*, and the reticulated networks lie towards larger *M* and smaller *D* (Ronel-lenfitsch et al. 2019). Whereas *ζ* parametrizes this Pareto front position as an input modeling parameter, it cannot be quantified from an observed network and can therefore not be used as a descriptor for predicting how the trade-off is balanced. We will use this formulation to generate networks along the Pareto front for which we then assess the relationship between these networks’ trade-off balance and their tree-to-cycle transition scale.

## 3 Tree-to-cycle transition scale as predictor of trade-off balance

Using Eq. (7) – (10) and *ζ* = 0.3 … 5.0, we generate networks (5 × 5 nodes) of different trade-offs of *D* and *M* . Details about the numerical implementation are provided in App. A. Figs. 1a and b show exemplary steady-state solutions of Eq. (10) with *β* = 2/3, *a* = 1, *b* = 1, and *c* = 0, i.e., without growth. The initial condition is a grid-like graph with edge conductivities randomly initialized in the range [0.5, 1.5]. This heterogeneity of initial conductivities pushes the network out of the artificial symmetry of the energy landscape.

**Figure 1:**
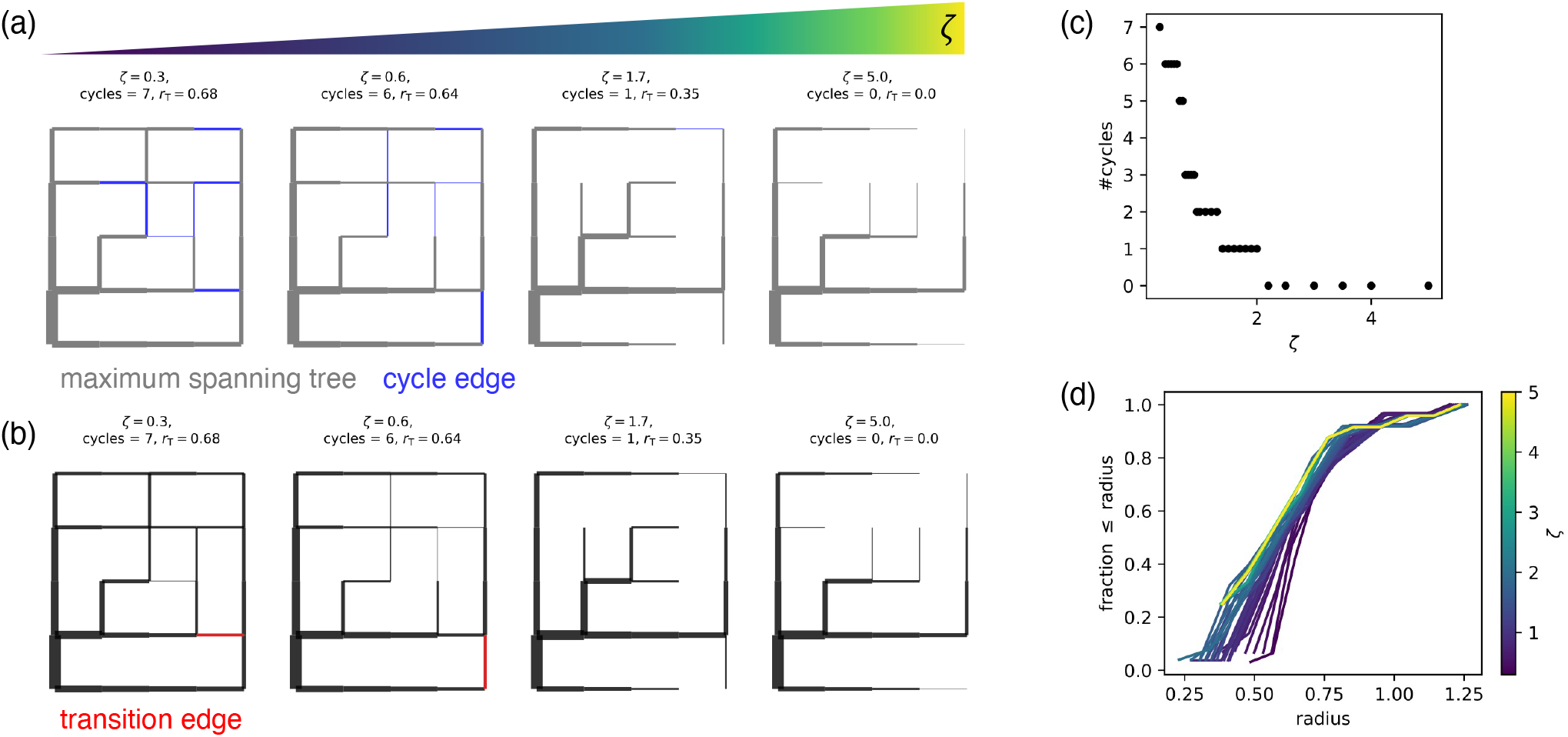
Analysis of adapted networks. (a,b) Steady-state networks whose reticulation degree decreases with increasing *ζ* (c) (in line with Ronellenfitsch et al. (2019)). After removing the maximum spanning tree (gray), the thickest of the remaining cycle edges (blue) is considered the transition edge (red). (d) The cumulative distribution of edge radii shows that the hierarchy is flatter the more reticulated the network. Adaptation parameters: *β* = 2*/*3, *a* = 1, *b* = 1, and *c* = 0.

### Graph analysis

We quantify the reticulation degree of these networks by computing the cyclomatic number, which is defined as (Barthélemy 2011):

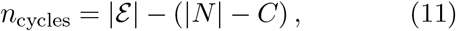

with the number of connected components *C* = 1. As shown in Fig. 1c, the cyclomatic number decreases with increasing *ζ* as fluctuations become more correlated and the relative difference in fluctuation between nodes decreases, aligning with previous results from Ronellenfitsch et al. (2019).

The cyclomatic number also influences the distribution of edge radii (Fig. 1a). To quantify this, we compute the cumulative distribution function (CDF) of the edge radii. The CDF characterizes the hierarchical structure of the graph: the more stretched the CDF, the wider the range of radius values, and the steeper the graph’s hierarchy. Conversely, if the CDF converges quickly, edges have more similar radii, and the hierarchy is flatter. Fig. 1d shows that the CDF converges more quickly for smaller *ζ*, which means that the more reticulated networks have a flatter hierarchy.

Using the hierarchy of edges of a given network, we define the graph’s tree-to-cycle transition scale as the radius of the thickest edge that does not belong to the graph’s maximum spanning tree (MST). Fig. 1a shows the MST in gray and Fig. 1b the transition edge in red. Intuitively, we interpret the thickness of this edge as a marker of the “maximal accepted cost” of forming a redundancy.

To enable quantitative comparison between graphs, we normalize *r*_T_ by the graph’s average edge radius, obtaining the dimensionless 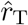. This is necessary because the absolute conductivity scale in Eq. (10) is set by the ratio *a/b* and is therefore arbitrary: two networks that differ by a uniform rescaling of all *k*_*ij*_ have the same architecture but different *r*_T_. Normalizing by the mean expresses the transition scale relative to a network’s own radius distribution, i.e., it locates the first cycle within the network’s hierarchy rather than on an absolute scale, and makes networks of different physical size directly comparable. Since the mean edge radius itself varies with *ζ* (Fig. 1d), we verified that the un-normalized *r*_T_ shows the same trends (Fig. S2), so the correlations reported below are not an artifact of the normalization. Hierarchical loop decompositions achieve a scale-independent characterization differently, by retaining the full sequence in which cycles merge under progressive removal of the weakest edges (Katifori et al. 2012; Modes et al. 2016); 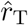 retains only the weight at which the first cycle appears, which is what makes it directly comparable across networks of different size and topology.

### The tree-to-cycle transition scale correlates linearly with the Pareto-front position

We compute 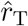 for steady-state networks for *ζ* = 0.3 … 5.0 without growth from five different initial conditions of randomly assigned edge conductivities. The trade-off plot of dissipation versus material cost of these networks (Fig. 2a) shows that 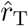 increases continuously along the Pareto front as the networks become more reticulated (color: inset legend). In particular, 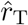 correlates negatively with *D* ^2^ and positively with *M* (see Fig. 2b and c, respectively). This means that the more the trade-off is shifted towards reducing *D* at the cost of increasing *M* , the larger 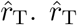 thus changes linearly along the trade-off balance.

**Figure 2:**
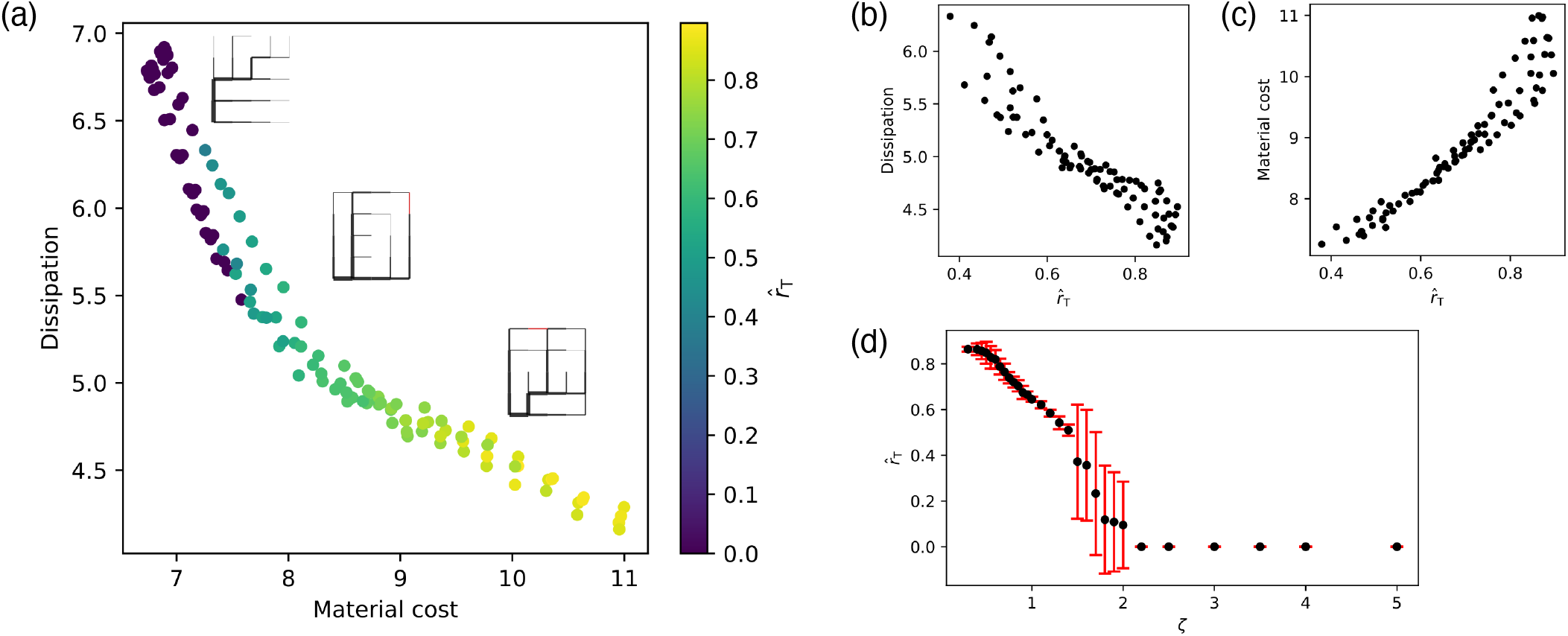
Tree-to-cycle transition scale correlates with Pareto-front position. By adapting networks to fluctuating flows with different correlation length scales *ζ*, Pareto-efficient networks with different degrees of reticulation are obtained. (a) Dissipation versus material cost of networks with different reticulation degree. Networks are steady-state solutions of the adaptation equation without growth for five initial conditions of a grid-like graph with edge conductivity randomly initialized in the range [0.5, 1.5]. 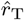 increases continuously along the Pareto front as the networks become more reticulated (color: inset legend). (b) 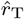 is negatively correlated with the dissipation. (c) 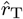 is positively correlated with the material cost. Data points without cycles 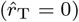 were excluded from (b) and (c). (d) 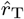 is negatively correlated with *ζ*. The standard deviation from the mean of 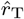 over the five samples (red error bars) is large for the transition from reticulated to tree-like structures, potentially because more networks are sub-Pareto-optimal in this region (see blue data points in (a)).

Another observation from Fig. 2b and c is that the data points deviate more strongly from the linear correlation towards larger *D* and larger *M* . We, therefore, asked whether the robustness of 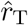 differs between locations in the Pareto space, i.e., network reticulation degrees. To analyze this, we compared the standard deviation of 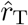 with respect to *ζ*. Fig. 2d shows that 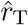 has a higher standard deviation for *ζ* between 1.3 and 2.0, which is the range at which the networks transit from cycled to tree-like.

Comparing this with Fig. 2a, we can find the same deviation in the upper left part of the Pareto front. In particular, only for some of the initial conditions did the networks converge to pure trees (dark purple points, 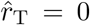); for the non-tree networks off the Pareto front, 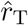 jumps up to ≈ 0.4. These sub-optimal networks likely arise from a local minimum in the energy landscape. This means that, in addition to the location on the Pareto front, 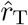 is sensitive to the distance from the Pareto front. This led to the question of how 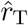 is related to the network’s adaptation trajectory through the energy landscape.

### The tree-to-cycle transition scale is sensitive to the adaptation trajectory

To test how the adaptation trajectory through the energy landscape affects 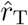, we add growth to the adaptation (Ronellenfitsch et al. 2016). For this, we repeat solving Eq. (10), but this time setting *c* = 1.0 and *λ* = 0.01, which corresponds to the parameter values that have been shown to produce networks close to the Pareto front (Ronellenfitsch et al. 2019). As before for the zero-growth case, we compute steady-state networks for *ζ* = 0.3 5.0 for five different initial conditions of randomly assigned edge conductivities.

The resulting data in Fig. 3a–d show that one effect of the growth term is that it evens out the effect of variations in the initial condition. In particular, whereas the steady-state networks without growth show visible variations across initial conditions, the data points of the networks with growth coincide, i.e., their standard deviation vanishes Fig. 3d.

**Figure 3:**
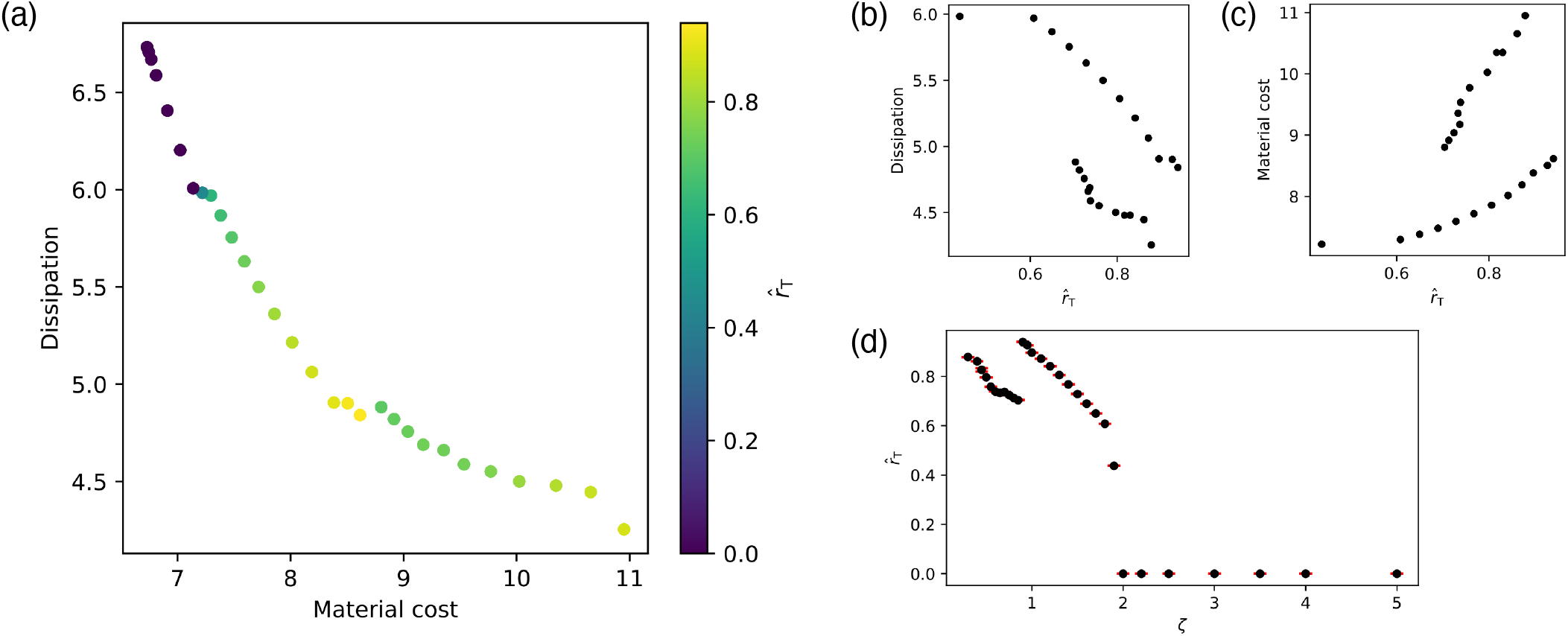
Adding a growth term to the adaptation leads to bifurcation of 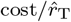 correlation. (a) Dissipation versus material cost of networks with different reticulation degree. 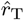 (color: inset legend) increases continuously along the Pareto front (decreasing *ζ*) until *ζ* = 0.85, then the value of 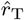 drops down before increasing again. This causes bifurcation in both the negative correlation of 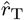 with the dissipation (b) and the positive correlation with the material cost (c). (d) 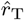 decreases as *ζ* increases until *ζ* = 0.85, then it jumps up before decreasing again. The growth term (*λ* = 0.01) cancels the effect of random initial conductivity (five samples) so that steady-state networks converge to the same 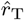, i.e., the standard deviation (red error bars) vanishes. Adaptation parameters: *β* = 2*/*3, *a* = 1, *b* = 1, and *c* = 1.

With respect to 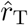, the growth term causes a bifurcation of the correlation between 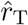 versus *D* and 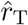 versus *M* (Fig. 3b and c, respectively). The point of bifurcation corresponds to a shift of networks away from the Pareto front for *ζ <* 0.85 (Fig. 3a). Growth is thus not uniformly beneficial across *ζ*: because it alters the adaptation trajectory, it can also route networks into a sub-optimal basin that the zero-growth dynamics avoids.

To better understand how minimizing *E* drives networks towards the Pareto front and how this changes 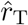 and its relationship with *D* and *M* , we track the corresponding values during the course of adaptation.

Fig. 4a shows that, without growth, networks shift gradually from the high *M* low *D* regime across space towards the Pareto front. During this process, 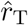, initially similar across networks, develops a gradient as the trade-off between *D* and *M* becomes increasingly different for different *ζ* (Fig. 4b and c, respectively). With growth, however, networks take a different optimization path across the multi-objective space (Fig. 4d), passing a much smaller subspace than without growth. The observation that with growth, networks are driven towards the Pareto front faster than without is in agreement with previous work (Ronellenfitsch et al. 2019). What is interesting here is that this progression, which is linear and smooth without growth, becomes disjointed with growth. This is reflected by jumps in 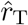 with respect to *D* and *M* as adaptation progresses (Fig. 4e and f, respectively). 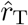 is thus sensitive to the adaptation trajectory.

**Figure 4:**
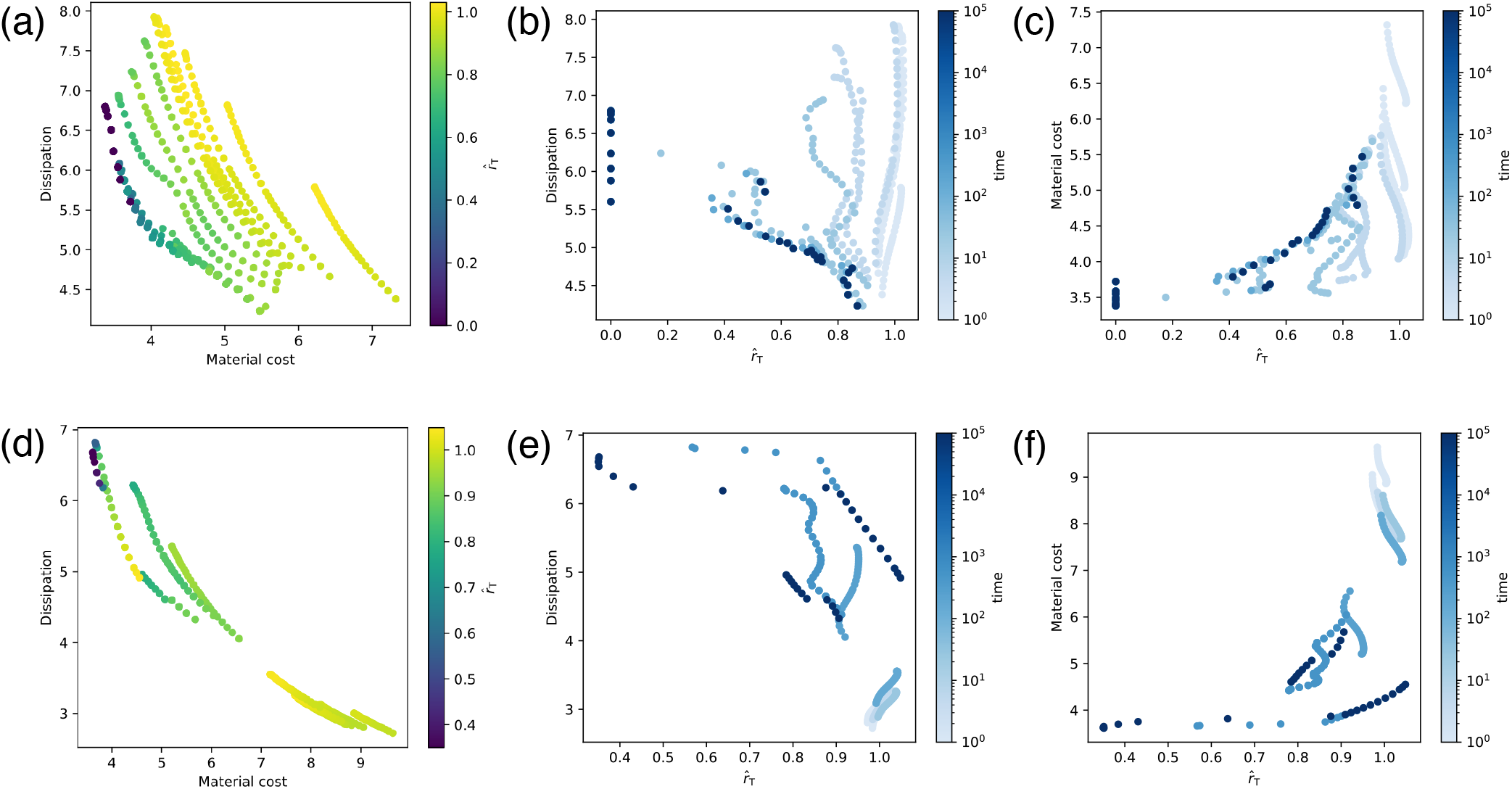
Gradient of 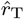 decreases as networks are driven closer towards Pareto optimality. Comparing adaptation without growth term (a, b, c) and with growth term (d, e, f) shows that networks are driven faster towards Pareto optimality by the growth term. However, at convergence of the adaptation without the growth term, more networks are tree-like (dark purple a versus d). Without the growth term, the slope of the correlation between 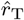 versus *D* (b) and 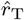 versus *M* (c) gradually increases. With the growth term (e, f), this shift is less smooth, and the correlation bifurcates, showing that 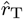 is sensitive to the adaptation trajectory. The time axis shows simulation time.

## 4 Conclusion

We have shown that the network’s tree-to-cycle transition scale characterizes how the trade-off between energy dissipation and material cost is balanced by the network. It can therefore be used as a descriptor for comparing how networks relative to each other will perform along these axes.

We further found that the tree-to-cycle transition scale is sensitive to the network’s adaptation trajectory and its distance from Pareto optimality. In particular, whereas close to the Pareto front, each network has a unique tree-to-cycle transition scale that correlates linearly with the position on the Pareto front, the transition scale of non-Pareto-optimal networks deviates from this correlation. The standard deviation of the transition scale across initial conditions is largest in the regime where some networks end up away from the Pareto front, and thus indicates how strongly the outcome depends on the adaptation trajectory.

When adding growth to the adaptation, the standard deviation of the transition scale vanishes. In this case, its correlation with the objective axis bifurcates for those modeling parameters for which networks end up away from the Pareto front.

Future directions include applying these descriptors to experimental data to advance our understanding of the principles governing biological adaptivity and resilience under varying environmental conditions.

## Acknowledgments

We thank Eleni Katifori for helpful discussions.

## Appendix A Implementation of dynamic vessel adaptation to correlated fluctuating flows

Based on Ronellenfitsch et al. (2019), we model dynamic vessel adaptation for correlated fluctuating sinks. For this, we impose boundary conditions with a source-sink tensor *S* = (***s***^(*m*)^) with dimensions |*N* | × |*N* |, where each column corresponds to a different microstate computed by Eq. (7). The source is node *l* = 0, located at the lower left corner of the lattice, and is held fixed across all microstates and all values of *ζ*. We evaluate the kernel elementwise on the |*N* | × |*N* | distance matrix *D*, whose entry *D*_*lm*_ = ∥***x***_*l*_ − ***x***_*m*_∥ is the Euclidean distance between nodes *l* and *m*, i.e., 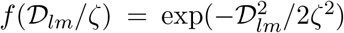; column *m* then holds the kernel centered at node *m*. Using this *S*, we compute an |*E*| × |*N* | dimensional flow matrix *Q* = (***q***^(*m*)^). From this *Q*, we obtain the |*E*| × 1 dimensional adaptation stimulus ⟨***q***^2^⟩ using Eq. (9) by averaging along each row of *Q* . We repeat this procedure for different values of *ζ*, thereby changing the width of the kernel function.

We compute Poiseuille flow rates ***q*** as follows^3^: (i) compute the |*N* | ×|*N* | weighted graph Laplacian from the oriented incidence matrix *B* and the diagonal conductance matrix *K* = diag(*k*_*ij*_*/l*_*ij*_), i.e., ℒ = *BKB*^T^, (ii) reduce system to (|*N* |−1)×(|*N* |−1) by removing first row and column of ℒ and corresponding entry in the source-sink vector, (iii) compute the pressure vector as 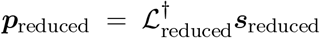, where † denotes the Moore-Penrose pseudo-inverse,^4^ (iv) recover the pressure at node *l* = 0: 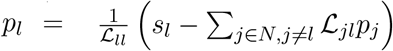, (v) compute the flow vector ***q*** = *KB*^T^***p***. Using ***q***, we then solve the dynamic vessel adaptation equation (Eq. (10)), using a fourth-order Runge-Kutta time-stepper and adaptation parameters *a* = *b* = 1 following previous work (Klemm et al. 2023). We stop when the relative total change 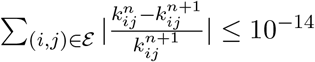.

## Supplementary figures

**Figure S1:**
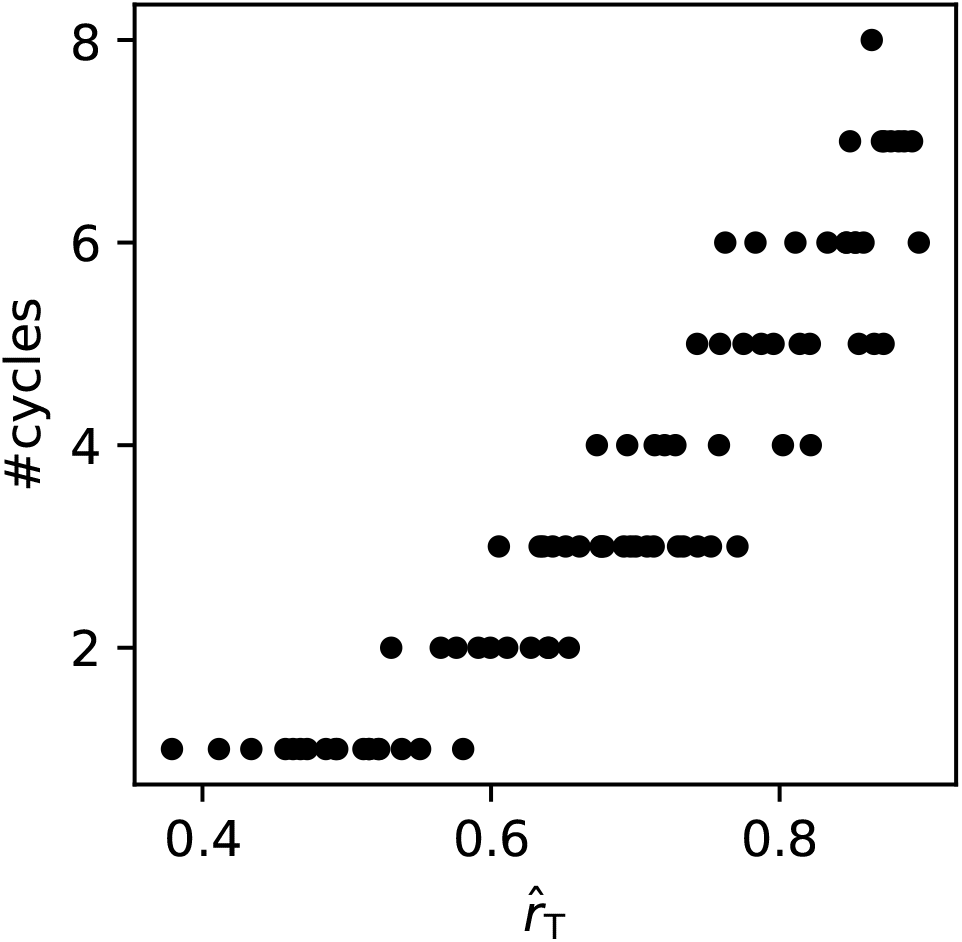
The cyclomatic number of the steady-state network is positively correlated with the tree-to-cycle transition scale. 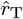 increases over a range of *ζ* at fixed cyclomatic number, then steps as a new cycle appears, producing a staircase rather than a smooth relationship. Adaptation parameters: *β* = 2*/*3, *a* = 1, *b* = 1, and *c* = 0.

**Figure S2:**
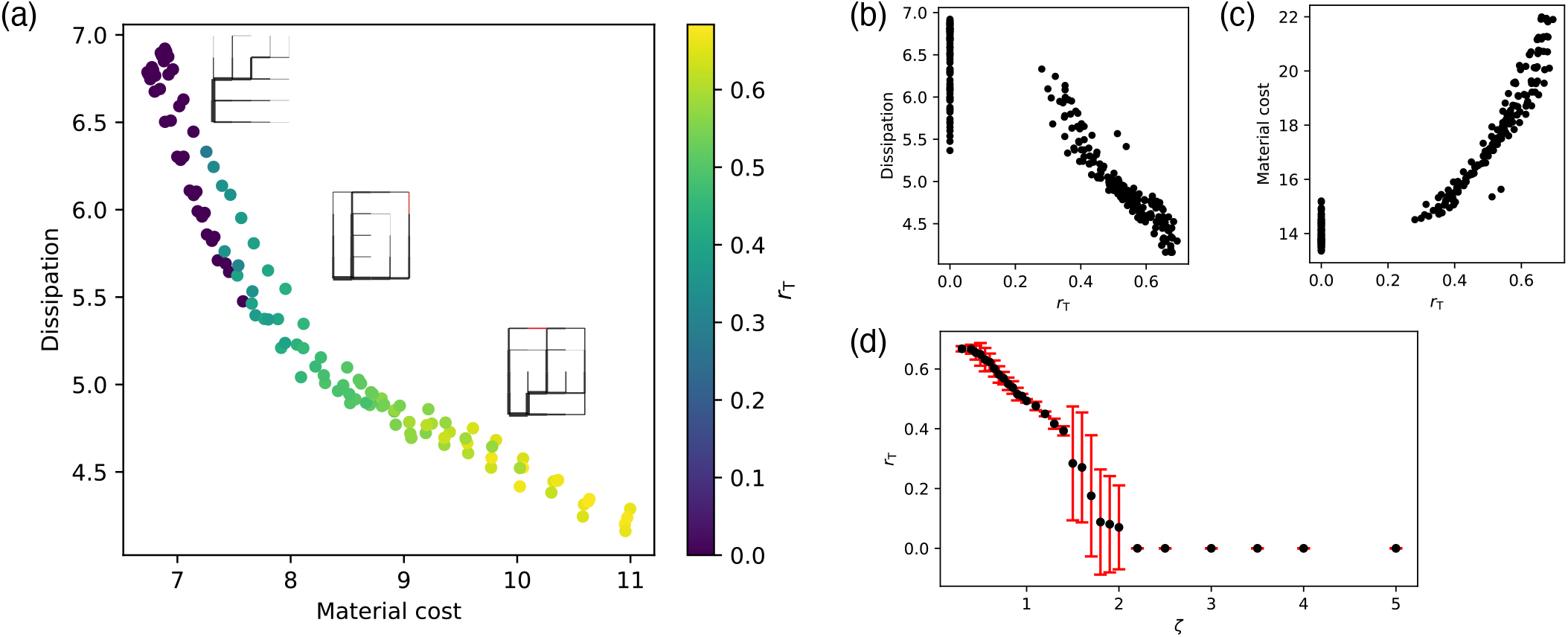
Unnormalized tree-to-cycle transition scale correlates with Pareto-front position similarly to the normalized transition scale. Panels (a)–(d) as in Fig. 2, but showing the unnormalized *r*_T_. All trends reported for 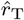 in the main text are reproduced, confirming that they are not an artifact of normalizing by the mean edge radius. Data points without cycles (*r*_T_ = 0) are here included in (b) and (c). Adaptation parameters: *β* = 2*/*3, *a* = 1, *b* = 1, and *c* = 0.

## Footnotes

1 second raw moment of the flow = mean flow plus the variance of the flow

2 *D* is computed for steady flow on the final network

3 Python libraries used: NetworkX, Numpy, Scipy

4 using scipy.linalg.pinv

